# Differential generation of saccade, fixation and image onset event-related potentials in the human mesial temporal lobe

**DOI:** 10.1101/442855

**Authors:** Chaim N. Katz, Kramay Patel, Omid Talakoub, David Groppe, Kari Hoffman, Taufik A. Valiante

**Affiliations:** Krembil Research Institute, Toronto Western Hospital (TWH), Ontario, M5T 1M8, Canada; Institute of Biomaterials and Biomedical Engineering, University of Toronto, Toronto, Ontario, M5S 3G9, Canada; Faculty of Medicine, University of Toronto, Toronto, ON, M5S 1A8, Canada; Perception & Plasticity Lab, York University, Toronto, ON, M3J 1P3, Canada; Department of Psychology, Vanderbilt University, Nashville, TN, 37240, United States of America; Division of Neurosurgery, Department of Surgery, University of Toronto, ON, M5S 1A1, Canada; Institute of Medical Sciences, University of Toronto, Toronto, ON, M5S 1A8, Canada; Electrical and Computer Engineering, University of Toronto, Toronto, ON, M5S 3G4, Canada

## Abstract

The electrophysiological signatures of encoding and retrieval recorded from mesial temporal lobe (MTL) structures are observed as event related potentials (ERPs) during visual memory tasks. The waveforms of the ERPs associated with the onset of visual stimuli (image-onset) and eye movements (saccades and fixations) provide insights into the mechanisms of their generation. We hypothesized that since eye movements and image-onset (common methods of stimulus presentation when testing memory) provide MTL structures with salient visual information, that perhaps they both engage similar neural mechanisms. To explore this question, we used intracranial electroencephalographic (iEEG) data from the MTLs of 11 patients with medically refractory epilepsy who participated in a visual search task. We sought to characterize electrophysiological responses of MTL structures to saccades, fixations and image onset. We demonstrate that the image-onset response is an evoked/additive response with a low-frequency power increase and post-stimulus phase clustering. In contrast, ERPs following eye movements appeared to arise from phase resetting of higher frequencies than the image onset ERP. Intriguingly, this reset was associated with saccade onset and not saccade termination (fixation), suggesting it is likely the MTL response to a corollary discharge, rather than a response to visual stimulation - in stark contrast to the image onset response. The distinct mechanistic underpinnings of these two ERP may help guide future development of visual memory tasks.

## INTRODUCTION

The electrophysiological responses of mesial temporal lobe (MTL) structures, known for their critical role in memory, have been characterized using a variety of behavioural paradigms and described according to their temporal relationship to stimulus onset (i.e. image-onset, word presentation, etc.). Neural responses following image presentation have been reliably used to study memory processes (Fell et al., 2004, 2008; Fernández et al., 1999; Montefusco-Siegmund et al., 2017; Paller and McCarthy, 2002; Sederberg et al., 2006; Tesche and Karhu, 2000), whereas, despite a rich literature linking eye movements to memory (for review see (Meister and Buffalo, 2016)), very few have investigated electrophysiological responses of human MTL structures to eye movements (Andrillon et al., 2015; Hoffman et al., 2013). At first glance, image-onset and saccadic eye movements both present salient information to visual centers, suggesting that subsequent processing would be similar, as would be the electrophysiological responses to them. Conversely, if such responses differ, it would suggest that their neural correlates differ, and that tasks/analyses that preferentially favor analyzing responses to one or the other are capturing distinct visual and likely mnemonic processes. Similarities and/or differences in these neural correlates could be representative of similarities and/or differences in the underlying neuronal mechanisms that generate these electrophysiological responses.

Arguments can be made to support or refute why MTL electrophysiological responses to image onset and saccadic eye movements might appear similar. Intuitively, since the hippocampus is a multimodal integrator far removed from primary visual cortex, it may be agnostic as to how visual information arrives (Rey et al., 2015). In this context any new retinal information, independent of its generation, would result in the same electrophysiological response. The literature however suggests visual and oculomotor responses are likely to be different. For example, MTL responses to the presentation of images, which are consistently used for investigations in word paradigms (Fell et al., 2008; Fernández et al., 1999), Sternberg probes (Tesche and Karhu, 2000), and complex visuals (Paller and McCarthy, 2002), are usually long latency responses with large amplitudes. (Andrillon et al., 2015; Mormann et al., 2005; Paller and McCarthy, 2002). On the other hand, eye-movement related responses in the MTL tend to have a shorter latency and lower amplitude (Andrillon et al., 2015). In line with these findings are those of Bartlett et al. who specifically compared the responses within macaque superior temporal sulcus (a region analogous to the human brain region responsible for integrating auditory and visual information (Beauchamp et al., 2004)) to image presentation and eye movements (Bartlett et al., 2011). Their results suggested a clear distinction between image-onset and eye-movement related responses in the time and frequency domain, and as well that the image-onset response can be modulated by succeeding eye movements. The precise origin of eye-movement related ERPs is likely dependent on where recordings have been performed, nevertheless there is indeed support for the concept that they in part represent a corollary discharge. Corollary discharges represent a transformed copy of a motor command signal that informs sensory systems of impending motor activity, thereby distinguishing self-generated changes in sensory signals from the changes generated from the external world (Crapse and Sommer, 2008). Such a signal can propagate widely throughout the brain, as well as through the ventral visual pathway and temporal lobe structures(Sommer and Wurtz, 2002) (Purpura et al., 2003). If indeed the MTL response to eye movements results in part from a corollary discharge, it would imply a fundamental mechanistic difference in how visual information is processed in mnemonic structures like the human hippocampus. However, it remains an open question whether the human mesial temporal lobe structures respond differently during these two processing conditions (i.e. image presentation and eye movements) – conditions that have substantial effects on memory and the corresponding electrophysiological measurements in the MTL (Andrillon et al., 2015; Axmacher et al., 2010; Fell et al., 2008; Hoffman et al., 2013; Kleen et al., 2016; Paller and McCarthy, 2002).

A variety of invasive and non-invasive electrophysiological recording techniques have been used to correlate MTL responses to underlying memory processes (Brewer et al., 1998; Jackson and Schacter, 2004; Long et al., 2014; McCormick et al., 2015; Merkow et al., 2015; Sederberg et al., 2006). Intracranial electroencephalography (iEEG), unlike other recording modalities, provides access to neural activity with high spatial and temporal resolution allowing investigation of electrophysiological activity in deep structures associated with memory processes, at ‘fast’ time scales (Johnson and Knight, 2015). Such activity can be seen in single trial events or averaged across trials aligned to events/stimuli to obtain event-related potentials (ERPs) (Davis, 1939). ERPs can be then further decomposed into their respective frequency components, to associate increases/decreases in power/phase clustering to encoding strength (Axmacher et al., 2010; Fell et al., 2008; Kleen et al., 2016; Paller and McCarthy, 2002) – so called subsequent memory effects (for review see (Hanslmayr and Staudigl, 2014)), and in the presence of ongoing oscillations, has been used to infer putative neuronal mechanisms like phase-resetting (Canavier, 2015; Fell et al., 2004; Hanslmayr et al., 2007; Klimesch et al., 2006). The study of ERPs has contributed significantly to our understanding of memory by allowing the attribution of changes in timing, amplitude, and spatial organization of brain related activity to putative cellular mechanisms thought to underlie the generation of brain oscillations (Womelsdorf et al., 2014).

ERPs are widely thought to be generated by either 1) stimulus-specific firing of additional neurons (evoked responses) or 2) post-stimulus phase-alignment of ongoing neural oscillations without additional neuronal firing (phase-reset response) (Makeig et al., 2002; Sauseng et al., 2007; Shah et al., 2004). Broadly speaking, the primary and most basic distinction between these two mechanisms is that evoked responses demonstrate an increase in individual trial power post-stimulus onset, whereas phase-reset responses do not (Shah et al., 2004). Identifying which of these mechanisms a specific ERP appears to be generated by not only characterizes the neuronal response, but also identifies the putative physiological processes that generate the response. For instance, in the auditory cortex of animals, ERPs arising from somatosensory stimulation appear as a phase-reset, arising from the “modulation” (Sherman and Guillery, 2002) of the timing of neuronal activity within the auditory cortex, without an increase in excitability (Schroeder and Lakatos, 2008). This preferential alteration of the timing of neuronal activity through phase-resetting is thought to be important in plasticity mechanisms, and transmission of information within the nervous system(Axmacher et al., 2006; Canavier, 2015; Voloh and Womelsdorf, 2016). Conversely, ERPs within the auditory cortex from auditory stimulation appear as an evoked-response, resulting from “driving” (Sherman and Guillery, 2002) increases in neuronal excitability (Schroeder and Lakatos, 2008). Therefore, the characterization of ERPs following image presentation and eye movements using this framework, can provide mechanistic insights to memory process in the human MTL. Furthermore, such a characterization provides a context to interpret existing memory work and guide the future design of tasks to probe human memory.

Thus to begin understanding the neural correlates of image onset and eye-movement associated ERPs in humans, MTL responses to image-onset and eye-movements were characterized and compared in epilepsy patients undergoing intracranial electroencephalographic (iEEG) investigations using a scene recognition task previously shown to have medial temporal lobe dependency (Chau et al., 2011). Intertrial phase clustering (ITPC) and spectral power were used to compare MTL electrophysiological responses to saccadic eye movements and image-onset. We demonstrate that the MTL responses following image-onset and saccade-onset events are starkly different with the former being a primarily evoked response showing within-trial power increases and phase clustering in lower frequency bands (delta and low theta) and the latter being best described as a phase-reset in the delta/theta/alpha frequency bands with significant phase clustering in the absence of within-trial power increases. We further demonstrate that saccade associated ERP are initiated with saccade

onset and not fixation, suggesting that the ERP associated with saccadic eye movements is initiated before retinal reafference, suggesting it is an internally generated signal like a corollary discharge (Andrillon et al., 2015; Purpura et al., 2003).

## RESULTS

### VISUAL SEARCH BEHAVIOUR

We used a previously reported behavioural paradigm (a change-blindness task) on a new set of participants that incorporates image onset and saccadic search, and has been shown to be MTL dependent (Chau et al., 2011) with neuronal correlates to search and memory localized to the hippocampus (Leonard et al., 2015, 2017; Montefusco-Siegmund et al., 2017). In this task, scenes were presented with the goal of finding the changing object in the scene. At the start of each trial, each participant was asked to fixate on a fixation cross in the middle of the screen, after which a scene, with a hidden target, was presented, subsequently referred to here as image-onset (see Star Methods for more details). All participants actively searched the scenes for the hidden target. Each participant contributed between 3000 and 21000 saccades/fixations across all experimental trials (see Table 1). Saccades had a median duration of 9ms and fixations had a median duration of 242ms (Figure 1B). Local field potentials were recorded from 15 hippocampal electrodes from seven different subjects and 16 parahippocampal electrodes from nine different subjects, ensuring that none of these electrodes were in the identified seizure-onset zone across all the patients (see Methods and Table 1 for more details).

**Figure 1.**
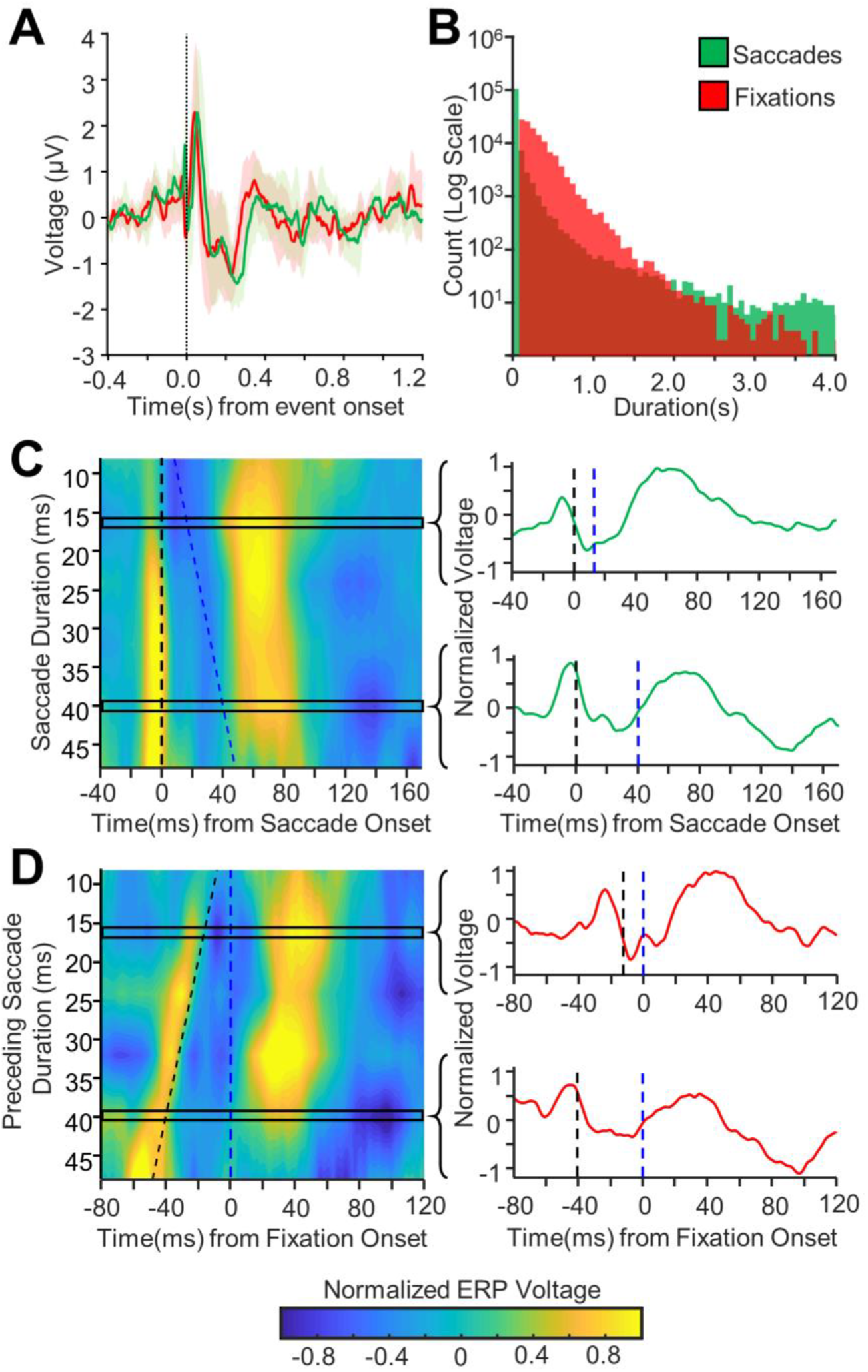
Saccade vs. fixation alignment of hippocampal response to eye movements. A: Grand-average Hippocampal ERP aligned to fixation (red) or saccade (green) onset (mean ± 95% confidence intervals are shown using corresponding colours). B: Histogram of fixation lengths (red) and saccade lengths (green) of all fixation lengths for all hippocampal electrodes. Y-axis has a log-scale to improve plot visibility. C: Left: Contour plot showing grand average saccade-aligned ERPs for changing saccade durations (shorter saccade durations towards the top of the graph, longer durations towards the bottom) for all hippocampal electrodes. The x-axis shows time relative to saccade onset. Saccade onset is marked by the dashed black vertical line (at time = 0ms). Fixation onset is marked by the dashed blue line. Note that as the saccade duration changes, the response remains aligned to the saccade onset. Right: the normalized ERP waveforms for two different rows of the contour plot is shown with the saccade onset marked with a dashed black line and fixation onset marked with the dashed blue line. Note, the saccade ERP is very similar for saccades of different durations across all electrodes. D: Left: As C but for fixation-aligned ERPs. Y-axis shows the duration of saccade preceding the fixation. The x-axis shows time relative to fixation onset. Fixation onset is marked with the dashed blue line (at time = 0ms), and saccade onset is marked with the dashed black line. Right: the normalized ERP waveforms for two different rows of the contour plot are shown with the saccade and fixation onsets marked as before. Note that as the saccade duration changes, the waveform shifts to remain aligned with saccade onset.

**Table 1.** Patient Summary Table. For each participant, the sex, resection zone, # of parahippocampal (PH) and # of hippocampal (HIPP) electrodes, # of saccades, median saccade duration, median fixation duration and # of experimental blocks completed are shown. Rows highlighted in black mark patients who were not analyzed due to insufficient data and poor eye-tracker calibration.

| ID | Sex | Age | Resection | PH | HIPP | # Saccades | Median Saccade Duration (ms) | Median Fixation Duration (ms) | # Blocks Completed |
| --- | --- | --- | --- | --- | --- | --- | --- | --- | --- |
| P14 | F | 28 | RAT | 2 | 0 | 12281 | 16 | 225 | 9 |
| P15 | M | 44 | RAT | 1 | 2 | 1909 | 75 | 109 | 2 |
| P16 | F | 39 | RAT | 2 | 4 | 16676 | 17 | 151 | 11 |
| P17 | F | 32 | NA | 1 | 2 | 3177 | 9 | 242 | 2 |
| P18 | M | 28 | NA | 1 | 1 | 17894 | 9 | 267 | 12 |
| P19 | M | 32 | NA | 4 | 1 | 4739 | 9 | 501 | 9 |
| P20 | M | 36 | NA | 1 | 2 | 14508 | 9 | 300 | 12 |
| P21 | F | 25 | RAT | 2 | 0 | 20717 | 16 | 217 | 12 |
| P22 | M | 31 | NA | 1 | 2 | 808 | 192 | 108 | 3 |
| P25 | F | 45 | RAT | 1 | 2 | 16701 | 9 | 276 | 12 |
| P26 | F | 58 | NA | 2 | 3 | 11267 | 9 | 259 | 6 |

**Table 2.**
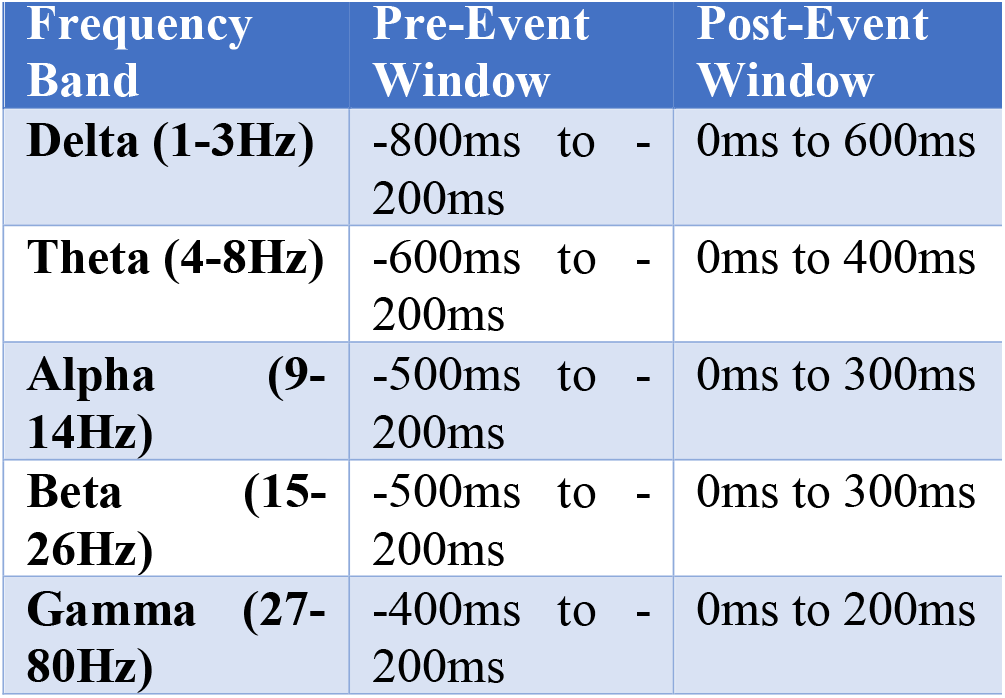
Time-frequency window definitions. This table shows the time-frequency windows that were used for the analyzing the changes in the phase clustering (ITPC) and power following image-onset and saccade-onset in this paper. Note that for each frequency band, the pre and post-event windows are of the same duration. Also, note that the pre-event windows were chosen to end 200ms before the onset of the event in an effort to avoid picking up any temporal smearing effects that may be present due to the short-time Fast Fourier Transform (stFFT) technique that was used to obtain the analytical signal.

| Frequency Band | Pre-Event Window | Post-Event Window |
| --- | --- | --- |
| **Delta (1-3Hz)** | -800ms to -200ms | 0ms to 600ms |
| **Theta (4-8Hz)** | -600ms to -200ms | 0ms to 400ms |
| **Alpha (9-14Hz)** | -500ms to -200ms | 0ms to 300ms |
| **Beta (15-26Hz)** | -500ms to -200ms | 0ms to 300ms |
| **Gamma (27-80Hz)** | -400ms to -200ms | 0ms to 200ms |

### FIXATION VS SACCADE RESPONSES

Since previous literature makes little distinction between eye-movement related responses in the MTL aligned to saccade-onset or to fixation-onset, these responses were analysed here for both alignments. Saccade-aligned and fixation-aligned responses in the MTL were nearly identical (Figure 1A). Saccade durations were very short (Figure 1B), suggesting that the saccade-onset ERPs and the fixation-onset ERPs were the same ERP aligned to two different, yet temporally proximal events. Although such short saccades are sometimes excluded from analysis or are considered to be microsaccades, they were included in the present analysis as they constituted a majority of the recorded saccades, and to ensure that the results reported here were generalizable across all saccades (Corrigan et al., 2018). To determine whether the observed response was better aligned to saccades or fixations, the saccade and fixation-onset ERPs were plotted as a function of saccade durations (Figure 1C and D, respectively). When saccade-onset ERPs (aligned to saccade onset) were plotted against changing saccade durations (Figure 1C), it was evident that the resulting ERPs remained aligned to saccade onset for varying saccade durations (i.e. vertical alignment of the contour plot in Figure 1C), suggesting that the response was aligned to saccade onset. To confirm this, the fixation-onset ERPs were plotted against preceding saccade durations (i.e. duration of the saccade preceding each fixation) in a plot aligned to fixation onset (Figure 1D). In this plot the observed fixation ERPs were still aligned to the saccade onset and not to the fixation onset (i.e. diagonal alignment of the contour plot in Figure 1D), further suggesting that the observed response was indeed a saccade-aligned response. Since the neural response to eye movements was determined to be aligned to saccade-onset, the saccade-onset ERPs were further analyzed using fixation-onset ERPs.

### ANALYSIS AND REMOVAL OF THE PERI-SACCADIC TRANSIENT

Saccade-onset ERPs across all analyzed electrodes contained a rapid transient response with a peak just prior to the onset of the saccade response (Figure 1A and Figure 2A/B left panel). To determine if this peak was an artifact resulting from volume conduction of muscle activity from the extraocular muscles, the spatial distribution of the peak-to-peak voltage of this transient was plotted across all the intracranial electrodes (see Figure S1 for a representative patient). These spatial distributions revealed that the transient was largest in the electrodes closest to the anterior temporal poles, which are also the ones closest to the orbit, providing strong evidence for this transient reflecting the potential associated with ballistic eye movements (electrooculogram - EOG) (See figure S2 for a spatial distribution of the artifact amplitude across all subjects compared to the spatial distribution of the saccade-onset response). Furthermore, previous work has shown that hippocampal recordings can contain eye movements artifacts, with waveforms and durations very similar to the transient observed in the current study (Kovach et al., 2011; Yuval-Greenberg et al., 2008). To test the alternate hypothesis that the transient was part of the physiological hippocampal response to eye movements, simulations were performed to determine whether a theta frequency phase reset mechanism (which has been previously reported as a potential mechanism through which the hippocampus responds to eye movements (Hoffman et al., 2013; Jutras et al., 2013)) can generate a high-frequency transient similar to the one that is seen in this study. The simulations demonstrated that a theta frequency reset mechanism can indeed generate a high-frequency transient depending on what phase the oscillators reset to (Figure 3A). However, the transient generated through this mechanism did not have the biphasic waveform observed in the transient identified in this study, suggesting that it is generated through a different mechanism. Interestingly, the simulated data showed broadband phase clustering around stimulus onset, despite being generated using a pure theta phase reset mechanism (Figure 3B). This was likely due to the high-frequency transient that was present in the simulated signals (Bénar et al., 2010; Kramer et al., 2008). This suggested that the high-frequency transient present in the recorded MTL response to eye movements would also skew the phase clustering metrics used for comparing the saccade-onset and the image-onset responses, further warranting its removal. Therefore, independent component analysis (ICA) along with a weighted average interpolation technique developed by Kovach and colleagues was used to remove this peri-saccadic transient from all the mesial temporal lobe recordings (Abel et al., 2016; Kovach et al., 2011). The removal of this transient substantially reduced peri-saccadic power in the 20-200Hz frequency band (in a 40ms window centred around saccade onset), which has been previously reported to be the frequency band in which a majority of the power of the intracranial EOG artifact resides (Kovach et al., 2011). The middle panels of Figure 2A and B show representative and grand average ERPs from hippocampal electrodes after this peri-saccadic transient was removed. Note that for all succeeding analysis of the saccade-onset response, the ICA-filtered data was used.

**Figure 2.**
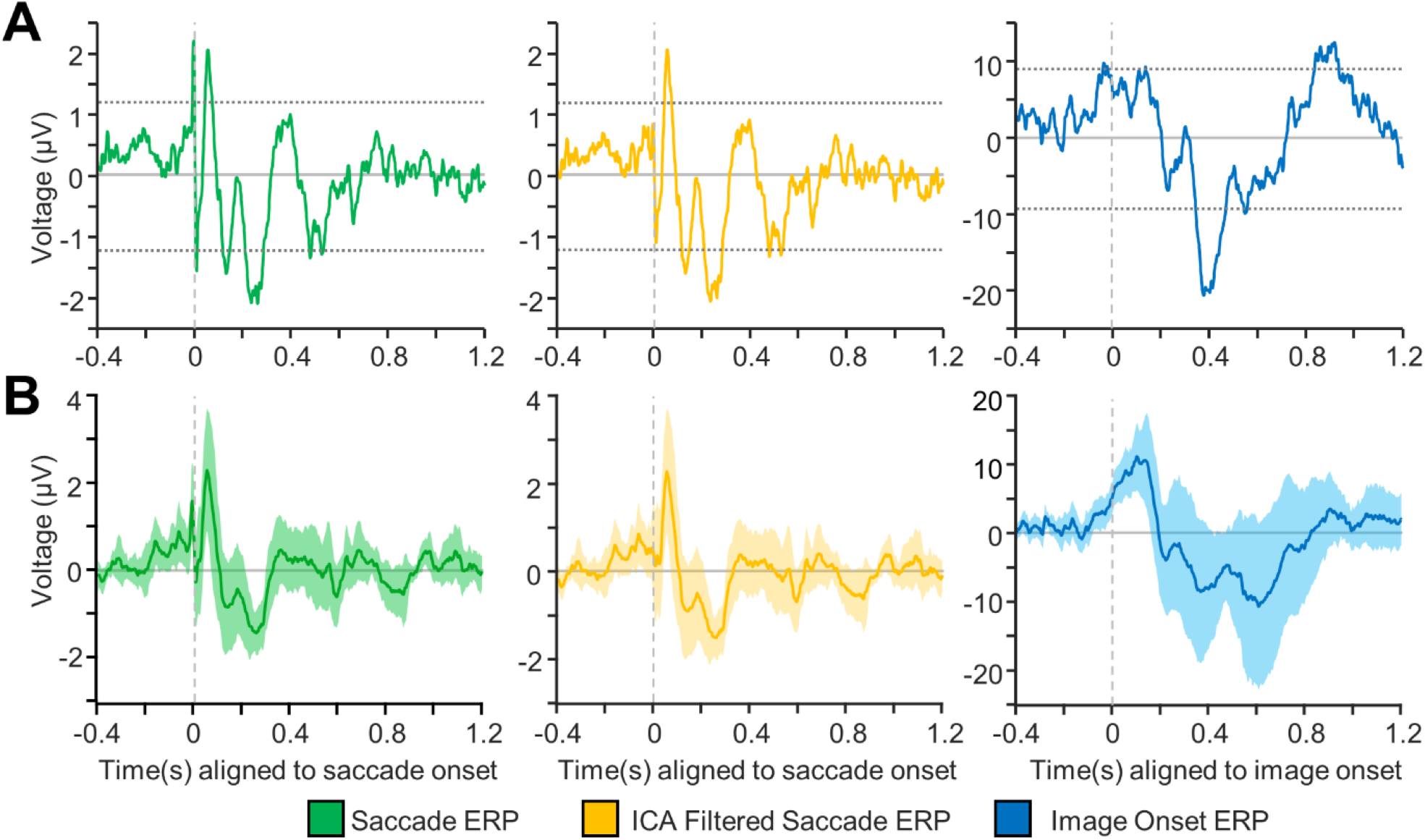
Event-related potentials (ERPs) following saccade and image onset. A: Representative ERPs from a hippocampal electrode for saccade aligned epochs (left/green), saccade aligned ICA-filtered epochs (middle/yellow) and image-onset aligned epochs (right/blue). The dashed horizontal lines mark significance bounds generated using permutation testing (3000 permutations) with randomized polarity inversions (see Method Details). B: Grand average ERPs for original saccade epochs (left/green), ICA-filtered saccade epochs (middle/yellow) and image-onset (right/blue) ERPs. The filled in region indicates 95% confidence intervals for each plot. Note that the ICA filtering successfully removes the transient, peri-saccadic response seen in the saccade aligned ERPs (green) while maintaining the integrity of the remaining ERP.

**Figure 3.**
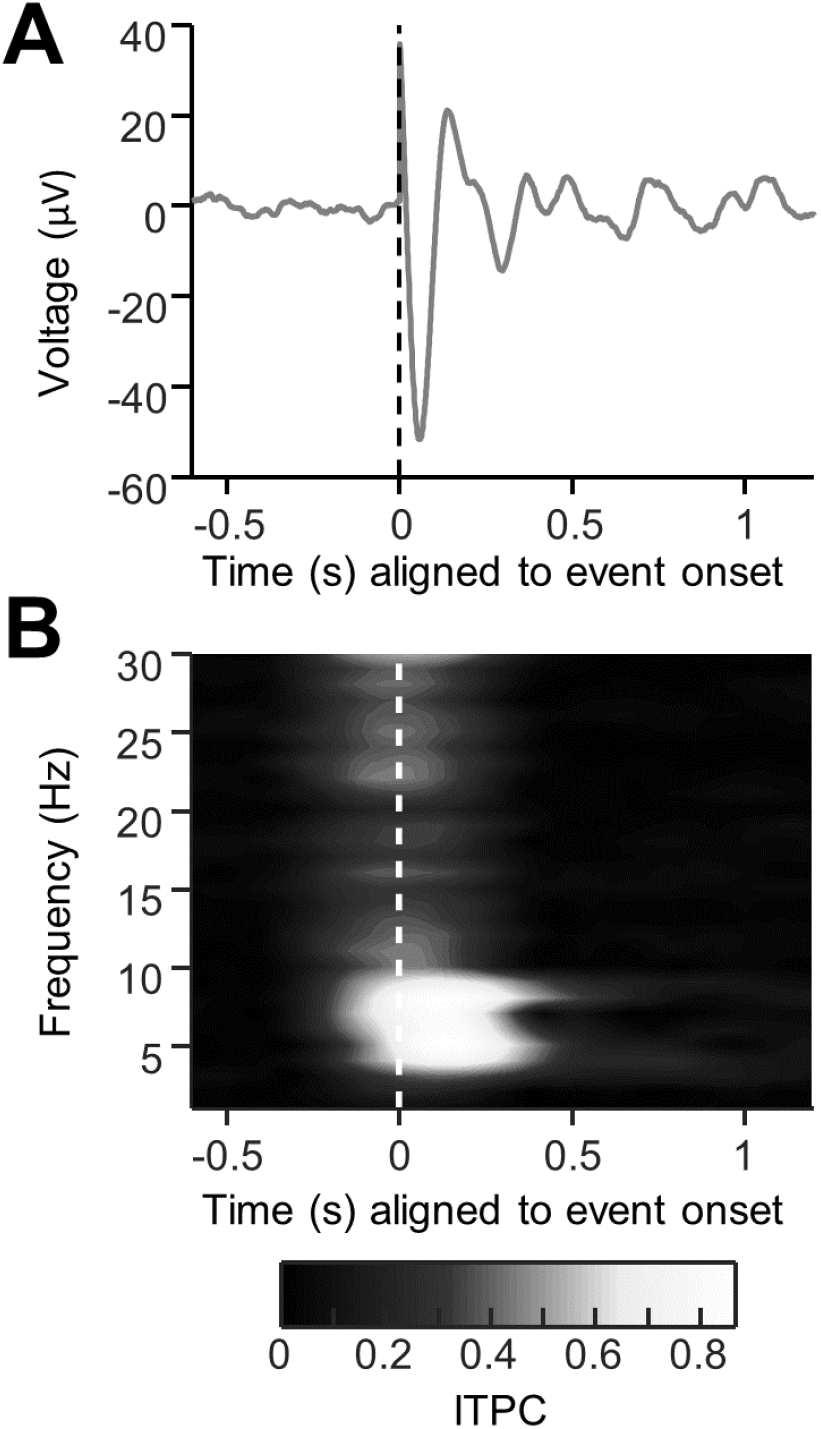
Theta phase reset simulation. A: An event-related potential generated by simulating a theta-band phase reset mechanism. To simulate this response, several oscillators (between 1 and 200 Hz) with random phase were summed together, and at time t=0, the phase of the oscillators in the theta frequency band was reset to a pre-selected phase (in this case, to 4/5 π or 144º). One thousand such simulated trials were summed together to generate this response. Notice the sharp, high frequency transient that is present in the ERP at time t=0 despite being generated using a low-frequency phase reset mechanism. B: The intertrial phase clustering of the same simulated data as in A. Notice that despite the mechanism of the response being a low frequency reset, broadband phase concentration is seen, suggesting that high frequency transients in the data can skew intertrial phase clustering metrics to show artificial, high frequency phase clustering that isn’t present in the underlying data.

### EVENT-RELATED POTENTIALS (ERPS)

To compare the temporal dynamics of the MTL response to saccade-onset and image-onset, event-related potentials (ERPs) were generated for both events. For each selected electrode, we calculated the average of the event-aligned response across all the trials, and across all experimental blocks. Non-parametric permutation testing was used to test the significance of this response. Figure 2A shows the ERPs for a representative hippocampal electrode for saccade epochs (left), ICA-filtered saccade epochs (middle) and for image onset epochs (right). 13 out of the 15 (86%) analyzed hippocampal electrodes had a statistically significant peak or trough following saccade onset and all hippocampal electrodes (100%) had a statistically significant peak or trough following image onset. Similarly, all parahippocampal (PHG) electrodes (100%) had a significant peak or trough following saccade onset and image-onset (Figure S6). The MTL response (hippocampal/PHG) to saccade-onset was more rapid compared to the image-onset response, which is evident from Figure 2. It is important to note that the observed ERPs had different amplitudes across subjects which is evident in other work (Paller and McCarthy, 2002) but may also have been influenced by the fact that the current analysis included all stimuli, without separating old and new items, and hits and misses, all of which have been shown the influence the amplitude of the image-onset response (Fell et al., 2004, 2008; Fernández et al., 1999; Montefusco-Siegmund et al., 2017; Mormann et al., 2005; Paller and McCarthy, 2002). To further characterize the difference in these responses, the phase clustering following stimulus onset was investigated.

### PHASE CLUSTERING

If, in the presence of an ongoing oscillation, phase clustering occurs following stimulus onset, then phase-resetting is considered the mechanism giving rise to the ERP. Hence, intertrial phase clustering (ITPC) was measured for saccade-aligned epochs and for image-onset aligned epochs. This measure assesses the distribution of the phase at each time point across trials and gives a value between 0 (no phase clustering, uniform phase distribution) and 1 (complete phase clustering). In this case, the ITPC value has been converted to an ITPCz value to control for the different number of trials for different electrodes/stimuli. The higher the value of the ITPCz, the higher the phase clustering. As evident from Figure 4, there is a stark contrast in the phase clustering following saccade and image-onset events. Significant hippocampal phase clustering was observed in the delta (1-3Hz), theta (4-8Hz) and alpha (9-14Hz) frequency bands. Significant hippocampal phase clustering was also observed following image-onset, however, it was largest in the delta frequency band and small, albeit significant, phase clustering was also observed in the theta and alpha frequency bands. The same analysis performed on all the parahippocampal electrodes (see Figure S6) revealed qualitatively similar results. It should be noted that the ITPC analysis reported here was performed on the ICA-filtered saccade-onset aligned epochs. When a similar analysis was performed on the original saccade-onset aligned epochs, broadband phase clustering was observed, likely due to the high-frequency peri-saccadic transient that was present in the original data. Although these results suggest significant phase clustering following image onset and saccade onset in the mesial temporal lobe structures, phase clustering can also be induced by an additive/evoked response that is added to ongoing oscillations at the onset of the stimuli (Lopour et al., 2013). To differentiate between this additive response and a true phase-reset response, the pre-and post-stimulus power was measured for individual trials and for the ERP.

**Figure 4.**
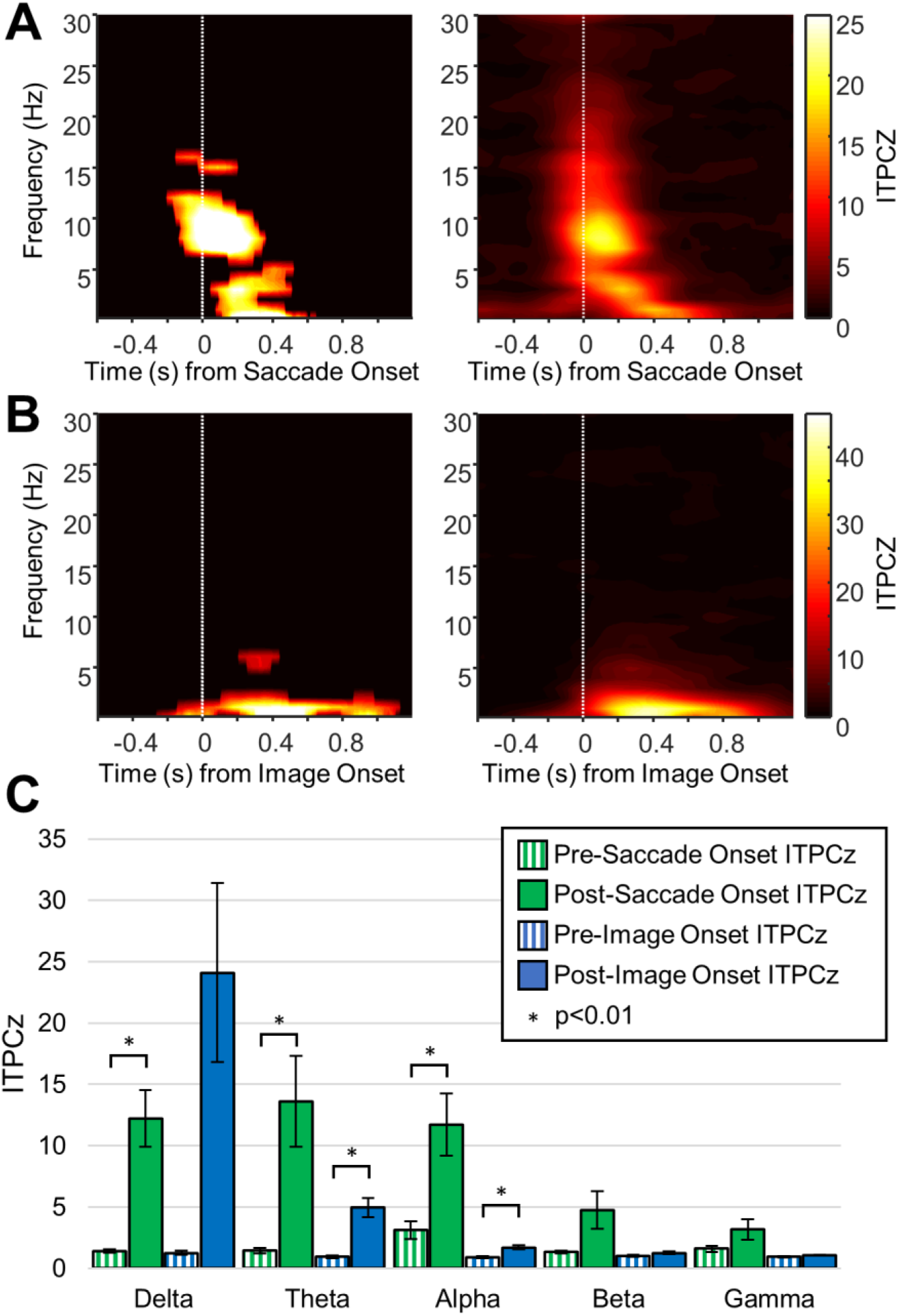
Intertrial phase clustering (ITPC) for hippocampal electrodes. A: Left-Representative ITPCz values (ITPC values that have been normalized to Rayleigh’s Z to control for a number of trials) for ICA-filtered saccade-aligned epochs. The plot has been masked to indicate significant values obtained using a Von-Mises distribution, which has been corrected for multiple comparisons using FDR correction at the α<0.001 significance level (see Method Details). Right – Grand average ITPCz value for ICA-filtered saccade aligned epochs for all hippocampal electrodes. Note the obvious phase clustering in the delta, theta and alpha frequency bands. B: Same as A but for image onset aligned epochs. Notice that the phase clustering is limited to lower frequency bands and has a longer time course. C: Comparison of ITPCz values for different time-frequency windows in the pre and post event periods (see Method Details for the definition of these time-frequency windows). Differences in the pre vs. post period were tested for significance using paired T-tests and corrected for multiple comparisons using FDR correction at the α<0.01 significance level. Significant differences are marked with an asterixis.

### POWER CHANGES

To investigate whether the observed phase clustering following saccade and image-onset events was due to a phase reset, the pre- vs post-event power was compared in the individual trials and in the ERP. For a pure phase-reset dependent ERP, there should be no power changes within each trial but there should be a significant increase in power following the event in the ERP. This is precisely what was observed for saccade responses (Figure 5A). There was no difference in spectral power in the individual trials across any of the frequency bands (Figure 5A-left and 5C), but there was a significant broadband increase in power (in the delta, theta, alpha, beta and low-gamma bands) following saccade onset in the saccade ERP (Figure 5A-right and 5D). Conversely, all the significant power changes in the image-onset ERP were mirrored by significant power changes in the individual trials (Figure 5B, C and D). This suggests that the image-onset response is not a characteristic phase reset, as there appears to be an addition of power to each individual trial following image onset, in contrast to the saccade response, which better fits the characteristics of phase resetting across the delta, theta and alpha frequency bands.

**Figure 5.**
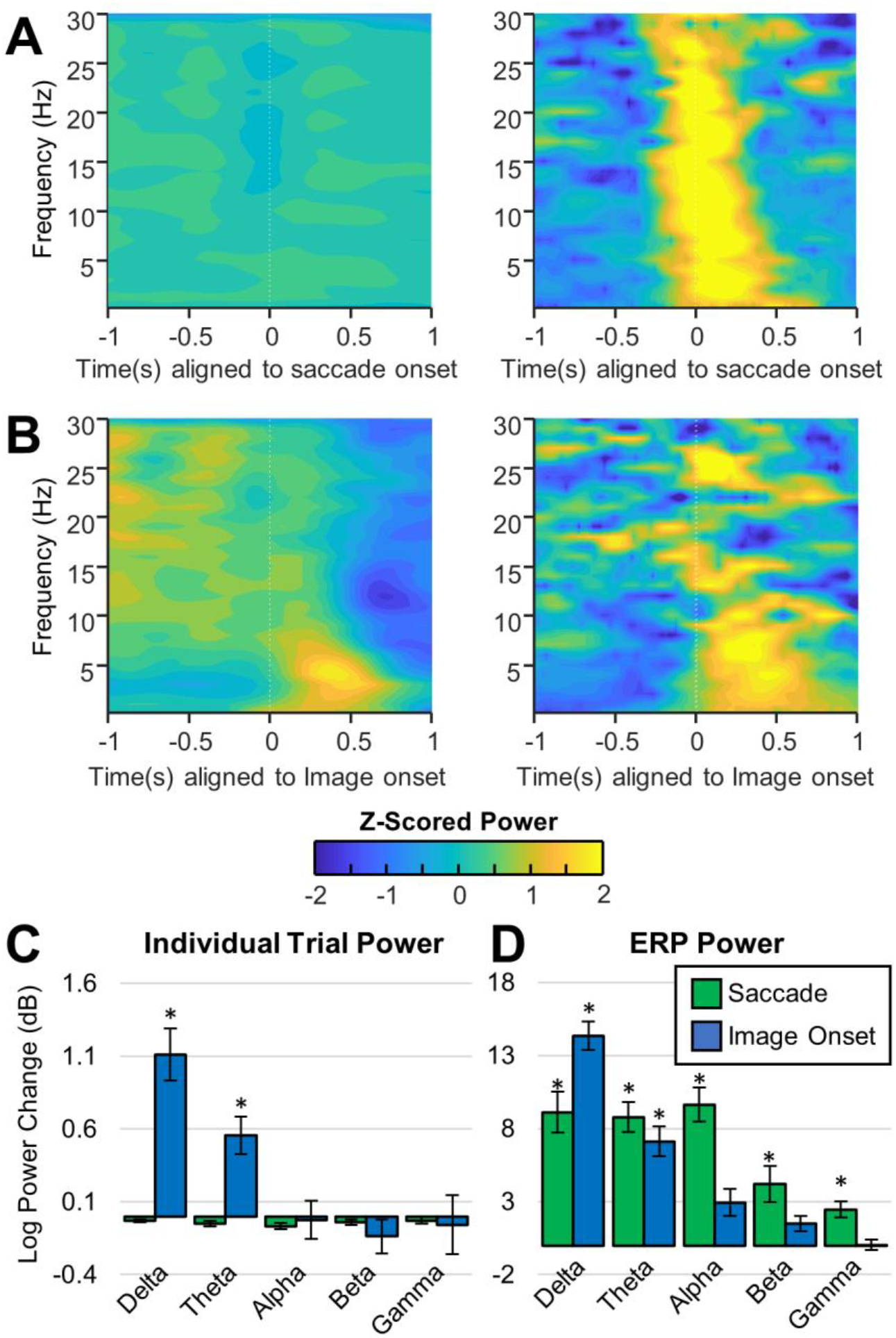
Spectral power for saccade-onset and image-onset epochs for hippocampal electrodes. A & B show power spectrograms in which the power for each frequency has been z-scored to better visualize power changes within each frequency. A: Left: Average Power of ICA-filtered individual saccade aligned epochs across all hippocampal electrodes. Right: Average power of ICA-filtered saccade-aligned ERPs averaged across all hippocampal electrodes. Notice that despite the significant increase in broadband power observed in the saccade ERPs, there is no observable increase in power in the individual trials. B: Left: Average power of individual image-onset aligned epochs across all hippocampal electrodes. Right: Power of image-onset aligned ERPs averaged across all hippocampal electrodes. Note that there is an observable increase in delta-band and theta-band power in the ERP and in the individual trials following image onset. C: Average log power change (post event power pre event power)) between pre and post event time-frequency windows (see Method Details for definition of the time frequency windows). Notice that there is no significant change in the power following saccade onset, but there is a significant increase in the power of the individual trials in the delta and theta bands following image onset. Significant changes in power are tested using a paired T-test, corrected for multiple comparisons using FDR correction. D: same as C but for the power of the ERP. Note that in the ERPs, there is a significant increase in broadband power following saccade onset and a significant increase in lower frequency power (delta and theta) following image onset.

## DISCUSSION

In this study, the intracranially recorded electrophysiological responses to image presentation and eye movements were investigated in the human MTL. These responses were characterized by investigating the underlying spectral characteristics of the corresponding event related potentials (ERPs). It was demonstrated that the MTL response following image presentation is a slow, evoked potential with corresponding phase clustering and power increases primarily in the delta band (Figure 4 and 5). In contrast, the MTL response following eye movements was a faster response with phase clustering but no evident within-trial power increases. This eye movement-response was observed to be aligned to the onset of the eye-movement (saccade) and not to the termination of the eye movement (fixation) (Figure 1). These findings suggest that there are distinct mechanisms through which these observed responses are generated, and the implications of these findings in the context of visual memory are discussed below.

## EVOKED IMAGE-ONSET RESPONSE

The image-onset ERP observed in this study was a slower potential than the saccade related ERP with a first trough occurring at 400-600ms, similar to the AMTL-N400 response that has been observed in the MTL and the PHG following the presentation of word stimuli (Fell et al., 2004; Fernández et al., 1999; Mormann et al., 2005, 2007) and had similar temporal characteristics as previously reported MTL response to visual stimuli (Paller and McCarthy, 2002) (Figure 2). Similarly, ERPs associated with probe presentation during a Sternberg task, demonstrated a late negative component, similar to this study but without the associated positive p300 component (Kleen et al., 2016).

The image-onset response observed in this study was accompanied by significant phase clustering in the theta and alpha bands. Additionally, the post-image onset phase-clustering was accompanied by an increase in theta and delta power (in individual trials and in the ERPs) with a latency of approximately 250-650ms. Together these observations suggest that the image-onset response is in large part an evoked response (Fell et al., 2004), and represents recruitment of additional neuronal populations (Mormann et al., 2007). This interpretation is consistent with observations that ERP amplitude, but not phase clustering, is decreased in hippocampal sclerosis (HS) (Mormann et al., 2007). Since the hallmark of HS is CA1 cell loss (Niemeyer, 1958), the reduction of the ERP amplitude in HS is likely directly related to the number of CA1 neurons that can be driven to threshold, as well as the number of synaptic contacts that can be made on them. The number of synaptic contacts will directly change the magnitude of the local field potential, as it is largely a manifestation of post-synaptic potentials (PSPs), which decrease in HS due to a reduction in the number of available synapses. Furthermore, since phase-clustering was not reduced (Mormann et al., 2005, 2007), a decrease in neuronal synchrony is unlikely to explain the decrease in ERP amplitude (Musall et al., 2014) in HS. At the cellular level it would be expected that an evoked response should be accompanied by an overall increase in cellular firing rates. Indeed studies have shown a marked firing rate increase at image onset peaking ~300ms after stimulus onset (Andrillon et al., 2015; Mormann et al., 2017), which is consistent with the latency of the image-onset ERP seen here(Figure 2). Our interpretations regarding the image onset ERP are consistent with a recent study that demonstrated an image-onset response very similar to the response observed here and characterized it as an additive/evoked response based on the amplitude/power and phase changes discussed earlier (Lopour et al., 2013). In summary, the image onset response that we observe appears to be a slower potential, with different frequency components than the saccade related ERP, and likely represents an evoked response (Lopour et al., 2013) that involves excitability increases of additional neurons.

## PHASE-RESET FOLLOWING SACCADE ONSET

Eye-movement related responses have been previously analyzed following saccade-onset (Jutras et al., 2013; Staudigl et al., 2017) and fixation (Hamamé et al., 2014; Hoffman et al., 2013; Rajkai et al., 2008). Here we were motivated by the work of Ito and colleagues who have shown that self-initiated saccades during the viewing of natural scenes were associated with phase resetting of the LFP within the beta frequency band, or in the theta band with a blank screen in area V1 of the macaque (Ito et al., 2011). Concomitant single unit recordings disclosed that the resetting was not associated with an increase in spike firing, but in fact was explained by a change in spike timing. They concluded that that the early (~75ms) LFP changes in V1 were likely the manifestation of eye movements associated corollary discharges (CD), whereas single unit firing rate changes were a manifestation of visual input. Similar findings have been demonstrated in area TE/STS (Bartlett et al., 2011; Purpura et al., 2003), and area V1 (Rajkai et al., 2008) in complete darkness, which suggest that the saccade related ERP is likely an modulatory input that phase resets ongoing oscillations.

In this context we have demonstrated that in the human MTL, the saccade associated ERP represents a phase-locking to saccades onset and not its termination (i.e. fixation) (Figure 1). Like others, we interpret these findings to imply that the post-saccade ERP arises not as a result of visual reafference, but a modulatory input like a corollary discharge that is transmitted from gaze control centers to mnemonic structures in addition to a number of other areas of the brain like FEF, TE, V1, and entorhinal cortex (EC) (Crapse and Sommer, 2008; Ito et al., 2011; Purpura et al., 2003; Sommer and Wurtz, 2002, 2008). Although it may be argued that our LFP analysis is too insensitive to definitively argue the ERP is comprised of a corollary discharge, prior work has shown that the saccade-associated LFP is reflective of eye-movements, whereas post-saccade firing rate changes reflect visual input (Ito et al., 2011).

Support for the speculation that a corollary discharge may be present in mnemonic structures comes from single unit studies in non-human primates (NHP), which have revealed the presence of saccade direction (SD) cells, some of which increase their firing rate as early as 68ms prior to the saccade (Killian et al., 2015). Previously, the same group reported saccade-related phase resetting of theta oscillations within non-human primate hippocampus during free-viewing, although no LFP data was presented in their more recent paper on SD cells (Killian et al., 2015). It would be interesting to understand if theta phase resetting occurs in the entorhinal cortex of the non-human primate, and the relationship of the SD cells to such ongoing population activity. Nonetheless, their hippocampal phase resetting effect was not associated with increases in spiking rate ((Jutras et al., 2013) Fig 1D), consistent with a modulating effect (as opposed to driving) (Sherman and Guillery, 2002) of a corollary discharge, as has been previously suggested in area TE of the NHP (Purpura et al., 2003). Such large-scale, cross-modal control via modulating inputs (phase resetting) has been suggested to be mediated through thalamic relay nuclei (Schroeder and Lakatos, 2008), which is further supported through anatomical connectivity between the memory and oculomotor networks (Shen et al., 2016). For entorhinal cortex SD cells, such a pathway via LD nucleus of the thalamus has been suggested as the route by which saccade direction information, possibly similar in content to the information transmitted to lateral inter-parietal sulcus (Duhamel et al., 1992), reaches the entorhinal cortex (Killian et al., 2015). Since the entorhinal cortex represents a major input to the hippocampus (Battaglia et al., 2011) hippocampal phase resetting associated with saccadic eye movements may arise from the transmission of SD cell information to area CA1 of the hippocampus.

That the saccade related phase-resetting starts with the onset of the saccade is strong evidence that corollary discharge (Ito et al., 2011; Purpura et al., 2003) underlies its generation, and argues against visual reafference, although additional characteristics of the response would be required to confirm its identity as a corollary discharge. Specifically, corollary discharges associated with saccadic eye-movements tend to be directional, and thus have different waveforms depending on the direction of the saccade (whether it is ipsiversive, or contraversive to the recording site). Furthermore, a saccade-related corollary discharge should be present in the dark. Although we have not shown this in our patients since obtaining complete darkness in the clinical setting is problematic, a number of studies have shown that both the LFP (Sobotka and Ringo, 1997), and single unit activity in the NHP hippocampus (Ringo et al., 1994) are modulated by saccades in complete darkness.

Although a number of studies in humans and NHPs have demonstrated both saccade related ERPs (Hoffman et al., 2013; Jutras et al., 2013), and alterations in single unit activity with saccadic eye movements (Bartlett et al., 2011), including those recording from the hippocampus, our analyses have allowed us to critically add to this literature by suggesting that the hippocampal ERP is associated with saccade onset, implying that it may arise from a corollary discharge (Ito et al., 2011; Purpura et al., 2003; Rajkai et al., 2008). What might be the role of such a corollary discharge within the hippocampus? Analogous to the discussion regarding SD cells in the entorhinal cortex (Killian et al., 2015), such saccade direction information may be useful for creating ‘mnemonic maps’ from visual stimuli, a process that requires knowledge of the viewing order, and spatial placement of the visual information. Additionally, corollary discharges in some regions of the brain provide the remapping transformation from the current receptive field (RF) of a neuron to the future field (FF) of the neuron (Duhamel et al., 1992). Envisaging the hippocampus as a comparator (Hasselmo, 2005; Vinogradova, 2008) of internal representation (via CA3 inputs) to current information (EC → CA1), and in the context of predictive-coding (Buckner, 2010), the remapping of current receptive fields to future fields in LIP (Duhamel et al., 1992), may by analogy represent the comparison of the future mnemonic representation (prediction) of the visual stimulus to the mnemonic representation of the current visual stimulus within the hippocampus.

This remains speculative, and future work will be required to understand the mechanistic role and information provided by such a phase-resetting/corollary discharge mechanism in the hippocampus beyond the temporal changes in excitability that accompany phase-resetting (Grover et al., 2009; Hyman et al., 2003; McCartney et al., 2004; Voloh and Womelsdorf, 2016)

Lastly it is important to note that saccade related phase-resetting was observed in other MTL sites such as the PHG which is known to play a role in visual search behavior (Supplementary Figures 2-5). For instance, parahippocampal damage has been previously shown to alter visual search behavior, leading to increased number of saccades and visual neglect (Mort et al., 2003; Nemanic et al., 2004). This is consistent with the previously cited literature that suggests anatomically widespread phase resetting associated with saccadic eye movements (Hoffman et al., 2013; Ito et al., 2011; Jutras et al., 2013; Purpura et al., 2003; Staudigl et al., 2017). This is not surprising if such saccade associated phase resetting is needed for preparing disparate brain regions for processing of new salient visual stimulus for information coding (Canavier, 2015), storage (Grover et al., 2009; Hyman et al., 2003; McCartney et al., 2004), and sensory integration (Voloh and Womelsdorf, 2016).

## COMPARISON OF THE TWO RESPONSES

The saccade-related ERP stands in stark contrast to the image-onset ERP that we observed, with the image onset response manifesting as an additive/evoked response, while the saccade response is better described as a phase resetting. At the cellular level it would be expected that an evoked response should likely demonstrate greater firing rate increases following image onset, which should occur later in time given its low frequency, whereas the saccade related ERP would likely not be accompanied by an increase in firing rate, would occur sooner after a saccade than the evoked response, and should show spiking phase-locked to the reset oscillation. Although there are limited studies to address this, a human single unit study by Andrillon et al (2016) comparing single unit image onset responses to saccade related responses demonstrated a 100% increase in firing rate after image onset, peaking at 300ms, while saccade related responses demonstrated an earlier increase in spiking, and a more modest firing rate increase of 25-30% following a saccade (Andrillon et al., 2015). Additionally, in the NHP hippocampus, an increase in firing rate is seen following image onset in the same task as the one used here, along with an evoked response and low-frequency phase clustering (Andrillon et al., 2015; Leonard et al., 2015; Montefusco-Siegmund et al., 2017), similar to the response reported in the current study. Although the human single unit data do suggests a modest increase in firing rate following a saccade, an excess of spike synchrony induced by a modulating input may underlie an increase in apparent firing rate (Ito et al., 2011) following saccades in humans (Andrillon et al., 2015) by biasing spike timing (Anastassiou et al., 2011; Cobb et al., 1995), and not increasing firing rate per se. Lastly and likely most importantly, the local field potential (LFP) is largely a manifestation of post-synaptic potentials, the extent of their synchronization (Buzsáki et al., 2012), and any associated spiking activity (Reimann et al., 2013). Recently it has been shown that the LFP is an excellent surrogate for predicting the contributions of post-synaptic currents to membrane potential fluctuations (Haider et al., 2016), and specific to our context, Ito and colleagues suggest that the early component of LFP following a saccade is an eye movement related potential and does not arise from visual reafference. Furthermore, cortical excitation has been observed following a saccadic eye movement in V1 in darkness, which likely arises from a post-inhibitory rebound (Rajkai et al., 2008; Ringo et al., 1994). Hence, we suggest that: 1) the LFP we observe following a saccadic eye movement has an extra-hippocampal origin that modulates the timing of hippocampal spiking but does not directly drive neurons to threshold; 2) it, like the responses in V1, are driven by an initial post-synaptic inhibition followed by rebound excitation (Rajkai et al., 2008; Ringo et al., 1994); 3) the rebound spiking acts non-locally (i.e.. projects to other brain regions) and thus itself does not contribute to the LFP recorded within the hippocampus and thus a power increase is not observed early after a saccadic eye movement and; 4) the image onset response reflects driving inputs from extra-hippocampal sites, likely entorhinal cortex, that are known to excite hippocampal neurons through glutamatergic synapses (Amaral and Witter, 1989; Battaglia et al., 2011; Suh et al., 2011; Yoshida et al., 2008). Future work will be required to elucidate the validity of these propositions.

## CONCLUSION

In this paper, the field potentials following saccade onset, fixation onset and image-onset were characterized using the temporal nature of the average MTL response, event-related phase clustering, and spectral power changes. Image-onset responses were shown to have low-frequency phase clustering and within-trial power increases, suggesting an additive/evoked neural response. In contrast, the saccade-related response was aligned to saccade onset, with significant phase clustering across the delta, theta, and alpha bands, and no within-trial power changes consistent with a phase-reset mechanism for the saccadic response. We propose that the saccade-onset response and image onset response are dichotomous, with the saccade-related response likely representing a corollary discharge that modulates hippocampal activity, whereas the image-onset response arises from driving sensory-related signals to the MTL. Future work will require further characterization of saccade related potentials at behavioral, and electrophysiological levels (LFPs and single units), specifically their generation in the dark, their directional dependencies, and the neuronal current sources in these regions. Our work has implications for memory-related testing and neuromodulation, since visual search is likely to engage different hippocampal mechanisms than would visual stimuli presented only during fixation.

## AUTHOR CONTRIBUTIONS

Conceptualization, C.K., and T.V.; Methodology, O.T. and K.H.; Software, C.K., O.T. and K.P.; Formal Analysis, K.P. and C.K.; Investigation, O.T., C.K. and K.P.; Data Curation, O.T., C.K. and K.P.; Visualization, K.P.; Writing – Original Draft, C.K. and K.P.; Writing – Reviewing and Editing, C.K., K.P., and T.V.; Funding Acquisition, K.H. and T.V.; Resources, T.V.; Supervision, K.H. and T.V.;

## ACKNOWLEDGEMENTS

We would like to acknowledge Victoria Barkley for her help with organizing and assisting with patient data collection. This work was supported by the Krembil Foundation, Brain Canada, NSERC, CIHR and Health Canada.

## CONFLICTS OF INTERESTS

The authors declare no conflicts of interest.

## Methods

### EXPERIMENTAL MODEL AND SUBJECT DETAILS

This study reports data from 11 subjects (6 females, 36.2±9.7 years of age) with medically refractory epilepsy, who were all implanted with subdural surface electrodes and depth macroelectrodes to localize epileptogenic regions (Table 1). Electrode locations were selected strictly on clinical considerations. These experiments were performed between 2 and 10 days post-operatively. All research was performed in accordance with protocols approved by the University Health Network Research Ethics Board.

### METHOD DETAILS

#### EXPERIMENTAL DESIGN

The experimental design of this study is identical to that described previously (Hoffman et al., 2013). A summary is provided here. Participants were seated comfortably in their hospital beds and a laptop computer was placed on a tray table in front of them, adjusted such that the screen was at approximately 60cm from their eyes. All participants first underwent a 9-point calibration of the eye-tracker system (iView RED – 120Hz, SensoMotoric Instruments, Teltwo, Brandenburg, Germany). The eye-tracker was connected to the laptop computer via a USB cable. Visual stimuli were presented on the laptop computer using Presentation (Neurobehavioral Systems, Albany, CA, USA). For each trial of the task, an original scene (taken from a large collection of natural scenes including landscapes, wildlife, cityscapes and indoor scenes) was shown at a full resolution of 1280×1024 and alternated with a target-modified scene every 500ms, with a 50ms grey screen separating these images. The target-modified scene was always a modified version of the original scene, in which one object in the scene (here called the ‘target’) was modified in Adobe Photoshop to give the impression that it disappeared. The size, location and content of these targets were modified between different scenes to reduce predictability. Participants were asked to search for the target in each pair of alternating images and could elicit the end of the trial by fixating their eyes on the target for a period of 1000ms. Once they found the target, or after a time limit of 45 seconds, the trial ended with revealing the target by rapidly altering between the two images without the grey screen gap. Trials were presented in blocks of 30, and each participant participated in up to 12 such blocks of data collection. Each scene pair was either novel or repeated once from a previous trial, with equal probability. Between trials in each block, participants were presented with a series of screens asking for verbal responses for the memory of the scenes and targets. Behavioural data from the inter-trial period is not presented or discussed here.

#### NEURAL RECORDINGS

Electrophysiological data presented here were recorded using depth macroelectrodes with four electrical contacts placed to record hippocampal activity. A 4-contact subgaleal electrode was used for ground and reference and was placed over the parietal midline facing away from the brain. Signals were sampled at 5kHz, and hardware filtered from 0.1 to 1kHz, with a NeuroScan SynAmps2 data acquisition system (Compumedics, Charlotte, NC, USA). Neural data was synchronized with eye-tracking data with the use of TTL triggers sent from the laptop computer to the NeuroScan computer. Electrode localization was performed by co-registering pre-op MRI with post-op CT using the iELVIS toolbox (Groppe et al., 2017). Following localization, the precise location of each of the 4 hippocampal electrodes was determined. Only those labelled as parahippocampal or proper hippocampal, and later verified by a neurosurgeon, were analyzed further.

#### DATA ANALYSIS

Eye tracking data was pre-processed to identify saccade and fixation events using the iView X iTools IDF Event Detector (SensoMotoric Instruments, Teltwo, Brandenburg, Germany). Fixation events were detected by this software using a dispersion based algorithm with a minimum fixation duration of 80ms and a maximum dispersion of 100 pixels (Salvucci and Goldberg, 2000). Saccade onsets were marked as the end of fixation events. All electrophysiological data was pre-processed in each trial by downsampling it at 1kHz, bandpass filtering between 0.5 and 200Hz using second-order Butterworth filter, and notch filtering at 30Hz and 60Hz to remove line noise and artifacts. All further analysis was performed using custom-written scripts in MATLAB (The Mathworks Inc, Natick, MA, USA).

##### Event-Related Potentials (ERPs)

To obtain ERPs for image-onset events, iEEG data from all trials and all relevant electrodes were aligned to the initial image-onset and trimmed to epochs containing 1.6 seconds of data before and after each event. The ERP for each electrode was then obtained by taking the mean of these image-onset epochs across all trials from all experimental blocks (1 image onset event per trial X 30 trials per blocks X up to 12 blocks per subject = ~360 image onset events for each ERP). Fixation and Saccade ERPs were obtained in a similar manner (±1.6ms around saccade and/or fixation onset) from the period following image onset, up until the end of the trial (when the target is found or revealed). As such, only saccades and fixations during the active visual search are used for analysis here. It should be noted that the temporal alignment of the ERPs is limited by the sampling frequency of the eye tracker (i.e. 120Hz) and the refresh rate of the display (i.e. 60Hz). This resolution may introduce some temporal jitter into the subsequent analysis, preventing elicitation of higher frequency phase clustering (if such were to exist).

##### Alignment of ERPs to fixation or saccade events

The extremely rapid nature of saccades (median length of 9ms across all subjects) makes it difficult to distinguish between a saccade-aligned ERP and a fixation-aligned ERP (Figure 1A). To determine whether the observed saccade-aligned ERPs and fixation-aligned ERPs were distinct events, or the same event viewed from different time points, the saccade-aligned and fixation-aligned ERPs were calculated for varying saccade durations. To do this, saccade and fixation epochs were binned based on the duration of the saccade succeeding the saccade-onset and preceding the fixation-onset. Considering the eye-tracker sampling rate of 120Hz, the first bin was centred at 8ms with a width of 8ms. Succeeding bins were separated by 8ms and had the same width. Saccade lengths greater than 56ms were not analyzed for the sake of this comparison because of the small number of saccades in each succeeding bin. Fixation and saccade-aligned ERPs were then obtained by averaging all the epochs within each bin. These ERPs were then normalized by z-scoring (to control for the varying number of epochs in each bin) and plotted on a contour plot (Figure 1C for saccade-aligned ERPs and Figure 1D for fixation-aligned ERPs), with varying saccade durations on the y-axis and time relative to saccade-onset (Figure 1C) or time relative to fixation-onset (Figure 1D) on the x-axis.

##### Saccadic Transient Analysis and Removal

The saccade ERPs across all subjects appeared to have a transient peak briefly preceding saccade onset, hereafter referred to as the peri-saccadic transient. A spatial distribution of the peak-to-peak amplitude of this transient across all the electrodes in the brain (analysis not shown here) demonstrated that the transient had the highest amplitude closest to the anterior temporal poles (closest to the orbits). This suggested that the transient response was a muscle artifact originating from the ocular muscles. To remove this transient from further analysis, independent component analysis was performed on all the mesial temporal lobe electrodes implanted in each patient (between 8 and 12 electrodes per patient) using the Bell-Sejnowski ICA algorithm (Bell and Sejnowski, 1995) implemented in the EEGLab toolbox for Matlab (Delorme and Makeig, 2004). The resulting components were used one at a time to reconstruct the original data, and the variance-accounted-for (VAF) between the reconstructed signal and the original signal was calculated for each of these reconstructions as shown in equation 1 –

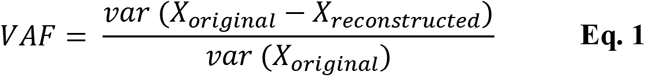

Here, *X_original_* is the original signal with the peri-saccadic spike, and *X_reconstructed_* is the signal reconstructed with each component, and *var* defines the variance of the signal. This VAF measure was only calculated for a 40ms window around each saccade event, to isolate the importance of each component towards generating the peri-saccadic transient. The VAF measure was then averaged for all saccades, and across all MTL electrodes for each reconstruction, and then ranked from largest to smallest for all the components. Based on this ranking, the two components with the largest contribution to the peri-saccadic transient were tagged for removal. Removing the entire component significantly affected the entire ERP waveform for the different electrodes. Thus, in an effort to preserve as much of the signal as possible, a moving average interpolation was performed on the two selected components, on a 40ms window around each saccade (±20ms)(Abel et al., 2016). The two modified components with the spikes removed were then used along with the non-modified components to reconstruct the original data, but with the peri-saccadic transient removed. This reconstructed data is referred to as the ICA-filtered data in all further analysis. Analysis of the ICA-filtered data demonstrated that the ICA-filtering significantly reduced the peri-saccadic power (±20ms around each saccade onset) in the 20-200Hz frequency band across all subjects, which has previously been reported as the frequency band in which a majority of the power of the EOG artifact is present in iEEG recordings (Kovach et al., 2011)

##### Phase Reset Simulations

To test the hypothesis that the peri-saccadic transient is part of the physiological hippocampal response to eye movements and not just an EOG artifact, simple simulations were performed to determine whether such a rapid transient response could be generated by physiological neural mechanisms. Prior work has suggested that the hippocampal response to eye movements (saccades or fixations) is generated by a phase reset or clustering mechanism, in which ongoing oscillators, in the theta frequency band, reset their phase following saccade or fixation onset. Hence, sinusoidal oscillators were simulated at every integer frequency from 1 to 200Hz with a randomly assigned phase. The amplitude of these oscillators was selected to match the power spectrum of the neural data recorded from a randomly selected hippocampal electrode. At time t=0, the phase of the oscillators in the theta frequency band was instantaneously reset to a predetermined phase, whereas the other oscillators continued to oscillate in accordance with their earlier phase. Temporal and frequency jitter was added to each oscillator to simulate the dynamics of real neural oscillators and to ensure that the phase alignment of the oscillators would quickly dissipate following the phase reset. The simulated neural signal “recorded” form each trial was the sum of the outputs of each oscillator. A thousand such trials were generated for each phase reset and an ERP was generated by summing the response across all of these trials. Figure 3 shows the ERP in response to the phase reset mechanism for a randomly selected phase. Intertrial phase clustering (ITPC) was also calculated for these simulated trials, as described below, and is also shown in Figure 3.

##### Intertrial Phase Clustering

Intertrial Phase Clustering (ITPC) is a measure used to look at how the phase of oscillations cluster in the time-frequency domain. To calculate ITPC, the time-frequency representation of all saccade and image-onset epochs was first obtained using a short-time Fast-Fourier Transform (stFFT). Note that the saccade-onset epochs were derived from the ICA-filtered data. A Hanning window of 1000ms, in steps of 10ms, was used to reduce edge artifacts. Equation 2 was then used to obtain the ITPC value. Here, *n* is the number of epochs (saccade or image-onset) for each electrode, and *θ_rtf_* is the phase angles of the *r^th^* epoch at time *t* and frequency *f*. The calculated value *ITPC_tf_* is the ITPC value at a given time-frequency point.

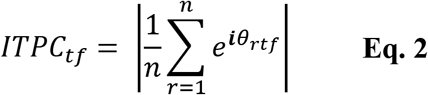

Since there were a significantly larger number of saccade epochs for each electrode compared to image-onset epochs, comparing the ITPC values between these events would have been difficult given the impact the number of events has on the calculated ITPC value. To reduce the effect of the number of events on the ITPC values, the values were transformed to ITPCz values, also known as Rayleigh’s Z, as shown in equation 3.

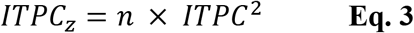

Average ITPCz values in five different, non-overlapping time-frequency windows (delta (1-3Hz), theta(4-8Hz), alpha(9-14Hz), beta(15-26Hz) and low-gamma(27-80Hz)) in the post and pre-event periods were calculated for further statistical analysis. The time duration of each of these windows in the post-event period was selected such that at least one full oscillation of the mean frequency component of each window were captured (see Table 2 for the precise temporal definition of these windows). Pre-event windows were chosen to be the same time duration as the post-event windows and were chosen to end 0.2 seconds before the event onset, to avoid picking up any post-event activity present before the event onset due to temporal smearing that may occur due to the selected Hanning window.

##### Pre- and Post-Stimulus Power

Spectral power was calculated for each individual epoch (saccade and image-onset) using the previously obtained time-frequency representation and then averaged across all trials for each electrode (note that the saccade-onset epochs were derived from the ICA-filtered data). Similarly, spectral power was also calculated for each electrode-specific ERP. To investigate power changes in each trial and in each ERP, pre-event and post-event power was obtained for the five different frequency bands defined in Table 2, by averaging out the power in the pre-event and post-event windows. Log power change (reported in decibels) between the post and pre-event windows was calculated using equation 4. Here, pre-event power refers to the average power in the pre-event time-frequency window and post-event power refers to the average power in the post-event time-frequency window –

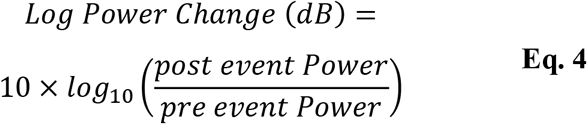

### QUANTIFICATION AND STATISTICAL ANALYSIS

#### REJECTED DATA

Experimental data was originally collected from 11 subjects, which contributed data from 19 hippocampi and 18 parahippocampi (Table 1). Note that data collected from hippocampi/parahippocampi that were later found to be in the seizure onset zone was not analyzed in this study and is not included in the counts provided in Table 1. Data from two of the subjects had to be rejected however due to poor eye-tracker calibration, incomplete experimental blocks and insufficient data. Hence, the data reported here is from 15 hippocampal electrodes from seven different subjects and 16 parahippocampal electrodes from nine different subjects. Note that for all population-level statistical analysis performed in this study, data points were marked as outliers using the Tukey method (Tukey, 1977) which is described as follows –

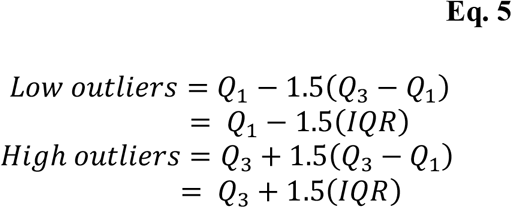

#### EVENT-RELATED POTENTIALS

To determine the significance of each ERP, a representative distribution of ERP maxima and minima was obtained for each electrode/ERP using non-parametric permutation testing with randomized polarity inversions (3000 permutations). ERP values were deemed significant if they fell above the 97.5^th^ percentile of the distribution of maxima, or below 2.5^th^ percentile of the distribution of minima.

#### INTERTRIAL PHASE CLUSTERING

##### Within-Electrode Analysis

Initial, within-electrode significance of each ITPCz value was obtained using equation 6, where *n* was the number of epochs used to obtain the ITPC values. This initial significance was then corrected for multiple comparisons using the Benjamini Hochberg FDR correction at the *α* < 0.001 significance level. Significant portions were then masked for visualization (Figure 3A)

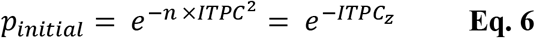

##### Between-Electrode Analysis

Between-electrode statistical analysis was performed to quantify the change in phase clustering in the post-event period compared to the pre-event period. To do this, specific time-frequency windows in the pre-onset and post-onset period for each event were defined, as shown in Table 2, and described earlier. The average ITPCz values from these windows were compared between all electrodes using a Paired, two-tailed T-test, corrected for multiple comparisons using the Benjamini Hochberg FDR correction at the *α* < 0.01 level.

#### SPECTRAL POWER ANALYSIS

The average power in the pre-and post-event period was obtained for five different frequency bands described earlier and shown in Table 2. The log power change was then calculated for each frequency band using equation 4. A two-tailed, one sample T-test was used to test the null hypothesis of zero power change following event onset. The significance values obtained using these tests were corrected for multiple comparisons using the Benjamini Hochberg FDR correction at the *α* < 0.01 significance level.

## REFERENCES

Abel, T.J., Rhone, A.E., Nourski, K. V., Ando, T.K., Oya, H., Kovach, C.K., Kawasaki, H., Howard, M.A., and Tranel, D. (2016). Beta modulation reflects name retrieval in the human anterior temporal lobe: an intracranial recording study. J. Neurophysiol. 115, 3052–3061.

Amaral, D.G., and Witter, M.P. (1989). The three-dimensional organization of the hippocampal formation: A review of anatomical data. Neuroscience 31, 571–591.

Anastassiou, C.A., Perin, R., Markram, H., and Koch, C. (2011). Ephaptic coupling of cortical neurons. Nat. Neurosci. 14, 217–224.

Andrillon, T., Nir, Y., Cirelli, C., Tononi, G., and Fried, I. (2015). Single-neuron activity and eye movements during human REM sleep and awake vision. Nat. Commun. 6, 7884.

Axmacher, N., Mormann, F., Fernández, G., Elger, C.E., and Fell, J. (2006). Memory formation by neuronal synchronization. Brain Res. Rev. 52, 170–182.

Axmacher, N., Cohen, M.X., Fell, J., Haupt, S., Dümpelmann, M., Elger, C.E., Schlaepfer, T.E., Lenartz, D., Sturm, V., Ranganath, C., et al. (2010). Intracranial EEG correlates of expectancy and memory formation in the human hippocampus and nucleus accumbens. Neuron 65, 541–549.

Bartlett, A.M., Ovaysikia, S., Logothetis, N.K., and Hoffman, K.L. (2011). Saccades during Object Viewing Modulate Oscillatory Phase in the Superior Temporal Sulcus. J. Neurosci. 31, 18423–18432.

Battaglia, F.P., Benchenane, K., Sirota, A., Pennartz, C.M. a, and Wiener, S.I. (2011). The hippocampus: Hub of brain network communication for memory. Trends Cogn. Sci. 15, 310–318.

Beauchamp, M.S., Lee, K.E., Argall, B.D., and Martin, A. (2004). Integration of Auditory and Visual Information about Objects in Superior Temporal Sulcus. Neuron 41, 809–823.

Bell, A.J., and Sejnowski, T.J. (1995). An Information-Maximization Approach to Blind Separation and Blind Deconvolution. Neural Comput. 7, 1129–1159.

Bénar, C.G., Chauvière, L., Bartolomei, F., and Wendling, F. (2010). Pitfalls of high-pass filtering for detecting epileptic oscillations: A technical note on “false” ripples. Clin. Neurophysiol. 121, 301–310.

Brewer, J.B., Zhao, Z., Desmond, J.E., Glover, G.H., and Gabrieli, J.D.E. (1998). Making memories: Brain activity that predicts how well visual experience will be remembered. Science 281, 1185–1187.

Buckner, R. (2010). The Role of the Hippocampus in Prediction and Imagination. Annu. Rev. Psychol. 61, 27–48.

Buzsáki, G., Anastassiou, C. a., and Koch, C. (2012). The origin of extracellular fields and currents — EEG, ECoG, LFP and spikes. Nat. Rev. Neurosci. 13, 407–420.

Canavier, C.C. (2015). Phase-resetting as a tool of information transmission. Curr. Opin. Neurobiol. 31, 206–213.

Chau, V.L., Murphy, E.F., Rosenbaum, R.S., Ryan, J.D., and Hoffman, K.L. (2011). A Flicker Change Detection Task Reveals Object-in-Scene Memory Across Species. Front. Behav. Neurosci. 5, 58.

Cobb, S.R., Buhl, E.H., Halasy, K., Paulsen, O., and Somogyi, P. (1995). Synchronization of neuronal activity in hippocampus by individual GABAergic interneurons. Nature 378, 75–78.

Corrigan, B.W., Gulli, R.A., Doucet, G., and Martinez-Trujillo, J.C. (2018). Characterizing Eye Movement Behaviors and Kinematics of Non-Human Primates During Virtual Navigation Tasks. J. Vis. 17, 1–22.

Crapse, T.B., and Sommer, M.A. (2008). Corollary discharge across the animal kingdom. Nat. Rev. Neurosci. 9, 587–600.

Davis, P. (1939). Effects of acoustic stimuli on the waking human brain. J. Neurophysiol. 2, 494–499.

Delorme, A., and Makeig, S. (2004). EEGLAB: An open source toolbox for analysis of single-trial EEG dynamics including independent component analysis. J. Neurosci. Methods 134, 9–21.

Duhamel, J.R.R., Colby, C.L., and Goldberg, M.E. (1992). The updating of the Representation of visual space in parietal cortex by intended eye movements. Science 255, 90–92.

Fell, J., Dietl, T., Grunwald, T., Kurthen, M., Klaver, P., Trautner, P., Schaller, C., Elger, C.E., and Fernández, G. (2004). Neural Bases of Cognitive ERPs: More than Phase Reset. J. Cogn. Neurosci. 16, 1595–1604.

Fell, J., Ludowig, E., Rosburg, T., Axmacher, N., and Elger, C.E. (2008). Phase-locking within human mediotemporal lobe predicts memory formation. Neuroimage 43, 410–419.

Fernández, G., Effern, A., Grunwald, T., Pezer, N., Lehnertz, K., Dümpelmann, M., Van Roost, D., and Elger, C.E. (1999). Real-time tracking of memory formation in the human rhinal cortex and hippocampus. Science 285, 1582–1585.

Groppe, D.M., Bickel, S., Dykstra, A.R., Wang, X., Mégevand, P., Mercier, M.R., Lado, F.A., Mehta, A.D., and Honey, C.J. (2017). iELVis: An open source MATLAB toolbox for localizing and visualizing human intracranial electrode data. J. Neurosci. Methods 281, 40–48.

Grover, L.M., Kim, E., Cooke, J.D., and Holmes, W.R. (2009). LTP in hippocampal area CA1 is induced by burst stimulation over a broad frequency range centered around delta. Learn. Mem. 16, 69–81.

Haider, B., Schulz, D.P.P.A., Häusser, M., and Carandini, M. (2016). Millisecond Coupling of Local Field Potentials to Synaptic Currents in the Awake Visual Cortex. Neuron 90, 35–42.

Hamamé, C.M., Vidal, J.R., Perrone-Bertolotti, M., Ossandón, T., Jerbi, K., Kahane, P., Bertrand, O., Lachaux, J.P., Hamamé, C.M., Vidal, J.R., et al. (2014). Functional selectivity in the human occipitotemporal cortex during natural vision: Evidence from combined intracranial EEG and eye-tracking. Neuroimage 95, 276–286.

Hanslmayr, S., and Staudigl, T. (2014). How brain oscillations form memories - A processing based perspective on oscillatory subsequent memory effects. Neuroimage 85, 648–655.

Hanslmayr, S., Klimesch, W., Sauseng, P., Gruber, W., Doppelmayr, M., Freunberger, R., Pecherstorfer, T., and Birbaumer, N. (2007). Alpha phase reset contributes to the generation of ERPs. Cereb. Cortex 17, 1–8.

Hasselmo, M.E. (2005). The Role of Hippocampal Regions CA3 and CA1 in Matching Entorhinal Input With Retrieval of Associations Between Objects and Context: Theoretical Comment on Lee et al. (2005). Behav. Neurosci. 119, 342–345.

Hoffman, K.L., Dragan, M.C., Leonard, T.K., Micheli, C., Montefusco-Siegmund, R., and Valiante, T.A. (2013). Saccades during visual exploration align hippocampal 3–8 Hz rhythms in human and non-human primates. Front. Syst. Neurosci. 7, 1–10.

Hyman, J.M., Wyble, B.P., Goyal, V., Rossi, C.A., and Hasselmo, M.E. (2003). Stimulation in hippocampal region CA1 in behaving rats yields long-term potentiation when delivered to the peak of theta and long-term depression when delivered to the trough. J. Neurosci. 23, 11725–11731.

Ito, J., Maldonado, P., Singer, W., Grun, S., and Grün, S. (2011). Saccade-Related Modulations of Neuronal Excitability Support Synchrony of Visually Elicited Spikes. Cereb. Cortex 21, 2482– 2497.

Jackson, O., and Schacter, D.L. (2004). Encoding activity in anterior medial temporal lobe supports subsequent associative recognition. Neuroimage 21, 456–462.

Johnson, E.L., and Knight, R.T. (2015). Intracranial recordings and human memory. Curr. Opin. Neurobiol. 31, 18–25.

Jutras, M.J., Fries, P., and Buffalo, E.A. (2013). Oscillatory activity in the monkey hippocampus during visual exploration and memory formation. Proc. Natl. Acad. Sci. 110, 13144–13149.

Killian, N.J., Potter, S.M., and Buffalo, E.A. (2015). Saccade direction encoding in the primate entorhinal cortex during visual exploration. Proc. Natl. Acad. Sci. 201417059.

Kleen, J.K., Testorf, M.E., Roberts, D.W., Scott, R.C., Jobst, B.J., Holmes, G.L., and Lenck-Santini, P.-P. (2016). Oscillation Phase Locking and Late ERP Components of Intracranial Hippocampal Recordings Correlate to Patient Performance in a Working Memory Task. Front. Hum. Neurosci. 10, 1–14.

Klimesch, W., Hanslmayr, S., Sauseng, P., and Gruber, W.R. (2006). Distinguishing the evoked response from phase reset: A comment to Mäkinen et al.. Neuroimage 29, 808–811.

Kovach, C.K., Tsuchiya, N., Kawasaki, H., Oya, H., Howard Iii, M.A., and Adolphs, R. (2011). Manifestation of ocular-muscle EMG contamination in human intracranial recordings. Neuroimage 54, 213–233.

Kramer, M.A., Tort, A.B.L., and Kopell, N.J. (2008). Sharp edge artifacts and spurious coupling in EEG frequency comodulation measures. J. Neurosci. Methods 170, 352–357.

Leonard, T.K., Mikkila, J.M., Eskandar, E.N., Gerrard, J.L., Kaping, D., Patel, S.R., Womelsdorf, T., and Hoffman, K.L. (2015). Sharp Wave Ripples during Visual Exploration in the Primate Hippocampus. J. Neurosci. 35, 14771–14782.

Leonard, T.K., Hoffman Correspondence, K.L., and Leonard, H. (2017). Sharp-Wave Ripples in Primates Are Enhanced near Remembered Visual Objects In Brief. Curr. Biol. 27, 257–262.

Long, N.M., Burke, J.F., and Kahana, M.J. (2014). Subsequent memory effect in intracranial and scalp EEG. Neuroimage 84, 488–494.

Lopour, B.A., Tavassoli, A., Fried, I., and Ringach, D.L. (2013). Coding of information in the phase of local field potentials within human medial temporal lobe. Neuron 79, 594–606.

Makeig, S., Westerfield, M., Jung, T.-P., Enghoff, S., Townsend, J., Courchesne, E., and Sejnowski, T.J. (2002). Dynamic brain sources of visual evoked responses. Science 295, 690–694.

McCartney, H., Johnson, A.D., Weil, Z.M., and Givens, B. (2004). Theta reset produces optimal conditions for long-term potentiation. Hippocampus 14, 684–687.

McCormick, C., St-Laurent, M., Ty, A., Valiante, T.A., and McAndrews, M.P. (2015). Functional and effective hippocampal-neocortical connectivity during construction and elaboration of autobiographical memory retrieval. Cereb. Cortex 25, 1297–1305.

Meister, M.L.R., and Buffalo, E.A. (2016). Neurobiology of Learning and Memory Getting directions from the hippocampus : The neural connection between looking and memory. Neurobiol. Learn. Mem. 134, 135–144.

Merkow, M.B., Burke, J.F., and Kahana, M.J. (2015). The human hippocampus contributes to both the recollection and familiarity components of recognition memory. Proc. Natl. Acad. Sci. 112, 14378–14383.

Montefusco-Siegmund, R., Leonard, T.K., and Hoffman, K.L. (2017). Hippocampal gamma-band Synchrony and pupillary responses index memory during visual search. Hippocampus 27, 425–434.

Mormann, F., Fell, J., Axmacher, N., Weber, B., Lehnertz, K., Elger, C.E., Fernandez, G., and Fernández, G. (2005). Phase/amplitude reset and theta-gamma interaction in the human medial temporal lobe during a continuous word recognition memory task. Hippocampus 15, 890–900.

Mormann, F., Fernández, G., Klaver, P., Weber, B., Elger, C.E., and Fell, J. (2007). Declarative memory formation in hippocampal sclerosis: an intracranial event-related potentials study. Neuroreport 18, 317–321.

Mormann, F., Kornblith, S., Cerf, M., Ison, M.J., Kraskov, A., Tran, M., Knieling, S., Quian Quiroga, R., Koch, C., and Fried, I. (2017). Scene-selective coding by single neurons in the human parahippocampal cortex. Proc. Natl. Acad. Sci. 114, 1153–1158.

Mort, D.J., Malhotra, P., Mannan, S.K., Rorden, C., Pambakian, A., Kennard, C., and Husain, M. (2003). The anatomy of visual neglect. Brain 126, 1986–1997.

Musall, S., Von Pfost, V., Rauch, A., Logothetis, N.K., Whittingstall, K., Von Pföstl, V., Rauch, A., Logothetis, N.K., and Whittingstall, K. (2014). Effects of neural synchrony on surface EEG. Cereb. Cortex 24, 1045–1053.

Nemanic, S., Alvarado, M.C., and Bachevalier, J. (2004). The Hippocampal/Parahippocampal Regions and Recognition Memory: Insights from Visual Paired Comparison versus Object-Delayed Nonmatching in Monkeys. J. Neurosci. 24, 2013– 2026.

Niemeyer, P. (1958). The transventricular amygdala-hippocampectomy in temporal lobe epilepsy. In Temporal Lobe Epilepsy, pp. 461– 482.

Paller, K.A., and McCarthy, G. (2002). Field potentials in the human hippocampus during the encoding and recognition of visual stimuli. Hippocampus 12, 415–420.

Purpura, K.P., Kalik, S.F., and Schiff, N.D. (2003). Analysis of Perisaccadic Field Potentials in the Occipitotemporal Pathway During Active Vision. J. Neurophysiol. 90, 3455–3478.

Rajkai, C., Lakatos, P., Chen, C.M., Pincze, Z., Karmos, G., and Schroeder, C.E. (2008). Transient cortical excitation at the onset of visual fixation. Cereb. Cortex 18, 200–209.

Reimann, M.W., Anastassiou, C.A., Perin, R., Hill, S.L., Markram, H., and Koch, C. (2013). A biophysically detailed model of neocortical local field potentials predicts the critical role of active membrane currents. Neuron 79, 375–390.

Rey, H.G., Ison, M.J., Pedreira, C., Valentin, A., Alarcon, G., Selway, R., Richardson, M.P., and Quian Quiroga, R. (2015). Single-cell recordings in the human medial temporal lobe. J. Anat. 227, 394–408.

Ringo, J.L., Sobotka, S., Diltz, M.D., and Bunce, C.M. (1994). Eye movements modulate activity in hippocampal, parahippocampal, and inferotemporal neurons. J. Neurophysiol. 71, 1285–1288.

Salvucci, D.D., and Goldberg, J.H. (2000). Identifying fixations and saccades in eye-tracking protocols. In Proceedings of the Symposium on Eye Tracking Research & Applications, pp. 71– 78.

Sauseng, P., Klimesch, W., Gruber, W.R., Hanslmayr, S., Freunberger, R., and Doppelmayr, M. (2007). Are event-related potential components generated by phase resetting of brain oscillations? A critical discussion. Neuroscience 146, 1435– 1444.

Schroeder, C.E., and Lakatos, P. (2008). Low-frequency neuronal oscillations as instruments of sensory selection. Trends Neurosci. 32, 9–18.

Sederberg, P.B., Schulze-Bonhage, A., Madsen, J.R., Bromfield, E.B., McCarthy, D.C., Brandt, A., Tully, M.S., and Kahana, M.J. (2006). Hippocampal and Neocortical Gamma Oscillations Predict Memory Formation in Humans. Cereb. Cortex 17, 1190–1196.

Shah, A.S., Bressler, S.L., Knuth, K.H., Ding, M., Mehta, A.D., Ulbert, I., and Schroeder, C.E. (2004). Neural Dynamics and the Fundamental Mechanisms of Event-related Brain Potentials. Cereb. Cortex 14, 476–483.

Shen, K., Bezgin, G., Selvam, R., McIntosh, Anthony, R., and Ryan, Jennifer, D. (2016). An Anatomical Interface between Memory and Oculomotor Systems. J. Cogn. Neurosci. 28, 1772–1783.

Sherman, S.M., and Guillery, R.W. (2002). The role of the thalamus in the flow of information to the cortex. Philos. Trans. R. Soc. B Biol. Sci. 357, 1695–1708.

Sobotka, S., and Ringo, J.L. (1997). Saccadic eye movements, even in darkness, generate event-related potentials recorded in medial sputum and medial temporal cortex. Brain Res. 756, 168–173.

Sommer, M.A., and Wurtz, R.H. (2002). A pathway in primate brain for internal monitoring of movements. Science 296, 1480–1482.

Sommer, M.A., and Wurtz, R.H. (2008). Brain Circuits for the Internal Monitoring of Movements. Annu. Rev. Neurosci. 31, 317–338.

Staudigl, T., Hartl, E., Noachtar, S., Doeller, C.F., and Jensen, O. (2017). Saccades phase-locked to alpha oscillations in the occipital and medial temporal lobe enhance memory encoding. PLoS Biol. 15, 1–15.

Suh, J., Rivest, A.J., Nakashiba, T., Tominaga, T., and Tonegawa, S. (2011). Entorhinal cortex layer III input to the hippocampus is crucial for temporal association memory. Science 334, 1415–1420.

Tesche, C.D., and Karhu, J. (2000). Theta oscillations index human hippocampal activation during a working memory task. Proc. Natl. Acad. Sci. 97, 919–924.

Tukey, J.W. (1977). Exploratory Data Analysis. Addison-Wesley 2, 688.

Vinogradova, O.S. (2008). Hippocampus as Comparator: Role of the Two Input and Two Output Systems of the Hippocampus in Selection and Registration of Information. 72, 1285–1286.

Voloh, B., and Womelsdorf, T. (2016). A Role of Phase-Resetting in Coordinating Large Scale Neural Networks During Attention and Goal-Directed Behavior. Front. Syst. Neurosci. 10, 1–19.

Womelsdorf, T., Valiante, T. a, Sahin, N.T., Miller, K.J., and Tiesinga, P. (2014). Dynamic circuit motifs underlying rhythmic gain control, gating and integration. Nat. Neurosci. 17, 1031–1039.

Yoshida, M., Fransén, E., and Hasselmo, M.E. (2008). mGluR-dependent persistent firing in entorhinal cortex layer III neurons. Eur. J. Neurosci. 28, 1116–1126.

Yuval-Greenberg, S., Tomer, O., Keren, A.S., Nelken, I., and Deouell, L.Y. (2008). Transient Induced Gamma-Band Response in EEG as a Manifestation of Miniature Saccades. Neuron 58, 429–441.

